# Non-destructive chemical sensing within bulk soil using 1000 biosensors per gram of matrix

**DOI:** 10.1101/2022.02.11.480126

**Authors:** Emily M. Fulk, Xiaodong Gao, Li Chieh Lu, Kelly R. Redeker, Caroline A. Masiello, Jonathan J. Silberg

## Abstract

Gene expression can be monitored in hard-to-image environmental materials using gas-reporting biosensors, but these outputs have only been applied in autoclaved matrices that are hydrated with rich medium. To better understand the compatibility of indicator gas reporting with environmental samples, we evaluated how matrix hydration affects the gas signal of an engineered microbe added to a sieved soil. A gas-reporting microbe presented a gas signal in a forest soil (Alfisol) when hydrated to an environmentally-relevant osmotic pressure. When the gas signal was concentrated prior to analysis, a biosensor titer of 10^3^ cells/gram soil produced a significant signal when soil was supplemented with halides. A signal was also observed without halide amendment, but a higher cell titer (10^6^ cells/gram soil) was required. A sugar-regulated gas biosensor was able to report with a similar level of sensitivity when added to an unsterilized soil matrix, illustrating how gas concentration enables biosensing within a soil containing environmental microbes. These results establish conditions where engineered microbes can report on gene expression in living environmental matrices with minimal perturbation, using biosensor titers that are orders of magnitude lower than the number of cells typically observed in a gram of soil.

## INTRODUCTION

Soil microbes live within spatially and temporally heterogeneous environments, where macroscale environmental properties, including climate and land management practices, can significantly alter microscale conditions^1,2^. The soil microbiome is responsible for many critical processes, such as controlling the formation and turnover of soil organic carbon, regulating nitrogen cycling, producing greenhouse gases, and forming essential plant symbioses^1–3^. However, the connection between macroscale environmental conditions, the microscale environment, and microbial behavior remains poorly understood^1–4^. Engineered microbes that function as biosensors provide a strategy to study how individual microbes respond to dynamically changing conditions in bulk environmental matrices like soils and sediments^4^. While early biosensors relied on fluorescent or ice nucleation protein outputs to monitor gene expression, these reporters require applications of high biosensor titers, sample processing, and extraction from matrices to enable quantification of the output signal^4–7^.

Volatile gas reporters of gene expression offer an opportunity to non-destructively monitor microbial behaviors in bulk matrices with minimal sample processing^4^. To date, methyl halides^8^, acetaldehyde^9,10^, hydrogen sulfide^11^, ethylene^12^, and methyl salicylate^13^ have all been leveraged as gas reporter outputs. Among these outputs, methyl halides have been used to the greatest extent in matrices. These outputs are generated by programming cells to conditionally express a methyl halide transferase (MHT)^8^, which catalyzes the production of CH_3_Cl, CH_3_Br, or CH_3_I from halide ions and S-adenosyl methionine^14^. When used as a metabolic output for biosensors, these chemicals can diffuse through soil matrices to the headspace of closed systems and be quantified using gas chromatography-mass spectroscopy (GC-MS) without disrupting the matrix. MHT reporters can produce a signal under both oxic and anoxic conditions, making them useful in materials containing oxygen gradients^8^. To date, MHT reporters have been used to study gene transfer in soil^8^, the effect of soil on the bioavailability of microbe-microbe signaling molecules^12^, and the effect of organic matter on the bioavailability of a microbe-plant signaling compound^15^. Additionally, MHTs have been used in synthetic soils to monitor the effects of different soil properties on the bioavailability of cell-cell signals^16^.

To obtain a robust indicator gas signal from environmental matrices, prior gas reporter studies have all used twice-autoclaved matrices that are hydrated with a nutrient-rich growth medium, which leads to osmolarities an order of magnitude above typical levels in the environment^8,12,16^. This growth medium has also been supplemented with halide salts at concentrations that are orders of magnitude higher than the levels found in natural matrices like soils^8,12,16,17^. Furthermore, biosensors have been added at titers that exceed the abundance of individual consortia members^18–20^ and are on par with the total cell population in environmental matrices^1,21,22^. When environmental matrices are modified in these ways, the structure and function of the native microbiome is disrupted, limiting the utility of gas biosensors to monitor native microbial consortia without perturbation^16,23–25^. To use biosensors under conditions that better reflect natural soil environments, there is a need to apply more sensitive detection methods to quantify trace indicator gas signals. To date, gas biosensor studies have used detection methods capable of quantifying CH_3_Br signals above ∼1,000 ppt (mol/mol) in the gas phase^12^, which is >100-fold higher than natural atmospheric levels (∼7 ppt, mol/mol)^26^. Gas concentration methods such as thermal desorption (TD) can be coupled with GC-MS to measure dynamic CH_3_X fluxes in the environment, but these methods have not yet been used to monitor gas outputs from engineered microbes^27–29^.

To better understand how environmental matrices impact indicator gas signals relative to the ambient background, we evaluated how matrix treatment affects detection of gas biosensor output from an Alfisol forest soil (Oxyaquic Glossudalf). Specifically, we assessed how the composition of the soil hydration medium, signal accumulation time, and matrix sterilization affected the indicator gas signal. We find that *Escherichia coli* biosensors present a robust signal when introduced into a sieved and homogenized Alfisol that was hydrated to environmentally-relevant osmolarity using nitrogen-limiting minimal growth medium, which reflects famine conditions observed in nature outside of hotspots and hot moments of microbial activity^1,2^. Additionally, we show that a signal can be obtained from a soil using natural halide levels present in the matrix. Amending the Alfisol with halides improves the indicator gas signal 1000-fold and enables detection of a sugar using as few as 10^3^ engineered cells/gram soil. This biosensor titer is orders of magnitude lower than both the total number of cells observed in a gram of soil (10^7^ to 10^12^ cells/gram soil)^1,21,22^ and the number of cells applied to enable previous gas biosensor studies (∼10^8^ cells/gram soil)^8,12^.

## RESULTS AND DISCUSSION

### Effect of halide amendments on gas signal

To obtain an environmental matrix for studying the gas reporter signal produced by an engineered microbe, we collected and characterized the A and B horizon of an Alfisol (Oxyaquic Glossudalf) from a forest near the San Jacinto River in Conroe, TX. This site has not experienced any agriculture since at least 1984. The physical and chemical properties for this soil are summarized in Table 1. This soil is nutrient-poor with a low organic matter content and halide concentrations that are below the estimated global averages for soil chloride (100 ppm), bromide (32 ppm), and iodide (3.9 ppm)^17^. Additional soil characterization included particle size distribution (Table S1), mineralogy (Table S2 and Figure S1), and water retention properties (Figure S2).

**Table 1.** Soil properties. Values are reported as averages (±1 standard deviation) from three technical replicates. *BDL, below detection limit.

|  | Texture | pH | Organic<br>C | Total N | Bulk<br>density | Cl<sup>-</sup> | Br<sup>-</sup> | I<sup>-</sup> |
| --- | --- | --- | --- | --- | --- | --- | --- | --- |
|  |  |  | (%) | (%) | gram/cm<sup>3</sup> | ppm | ppm | ppm |
| A (0-10 cm) | Sandy<br>loam | 7.0 | 4.43 | 0.20 | 1.34 ($\pm 0.09$) | 16.2 ($\pm 0.6$) | 1.8 ($\pm 0.2$) | BDL* |
| B (10-58 cm) | Loamy<br>sand | 6.8 | 0.40 | BDL* | 1.87 ($\pm 0.09$) | 8.6 ($\pm 0.9$) | 0.9 ($\pm 0.6$) | BDL* |

Initially, we evaluated the gas signal from an *E. coli* strain that constitutively expresses an MHT (MG1655-*mht*)^8^ within a twice-autoclaved B horizon soil. We hydrated the soil to 32% field capacity with M63 medium containing 10^7^ colony forming units (CFU) per gram of dry soil and a range of NaCl or NaBr concentrations (Figure 1a-b). Following a 24-hour incubation, we observed a significant CH_3_Cl signal in soil samples having ≥10 mM supplemental NaCl (p < 0.01, two-sided *t* test). While the average CH_3_Br signal was higher than the detection limit in the absence of supplemental NaBr, ≥10 mM NaBr was also required to yield a significant signal (p < 0.01, two-sided *t* test). Negative controls performed using soil augmented with wildtype *E. coli* MG1655 did not present detectable CH_3_X under any condition tested.

**Figure 1.**
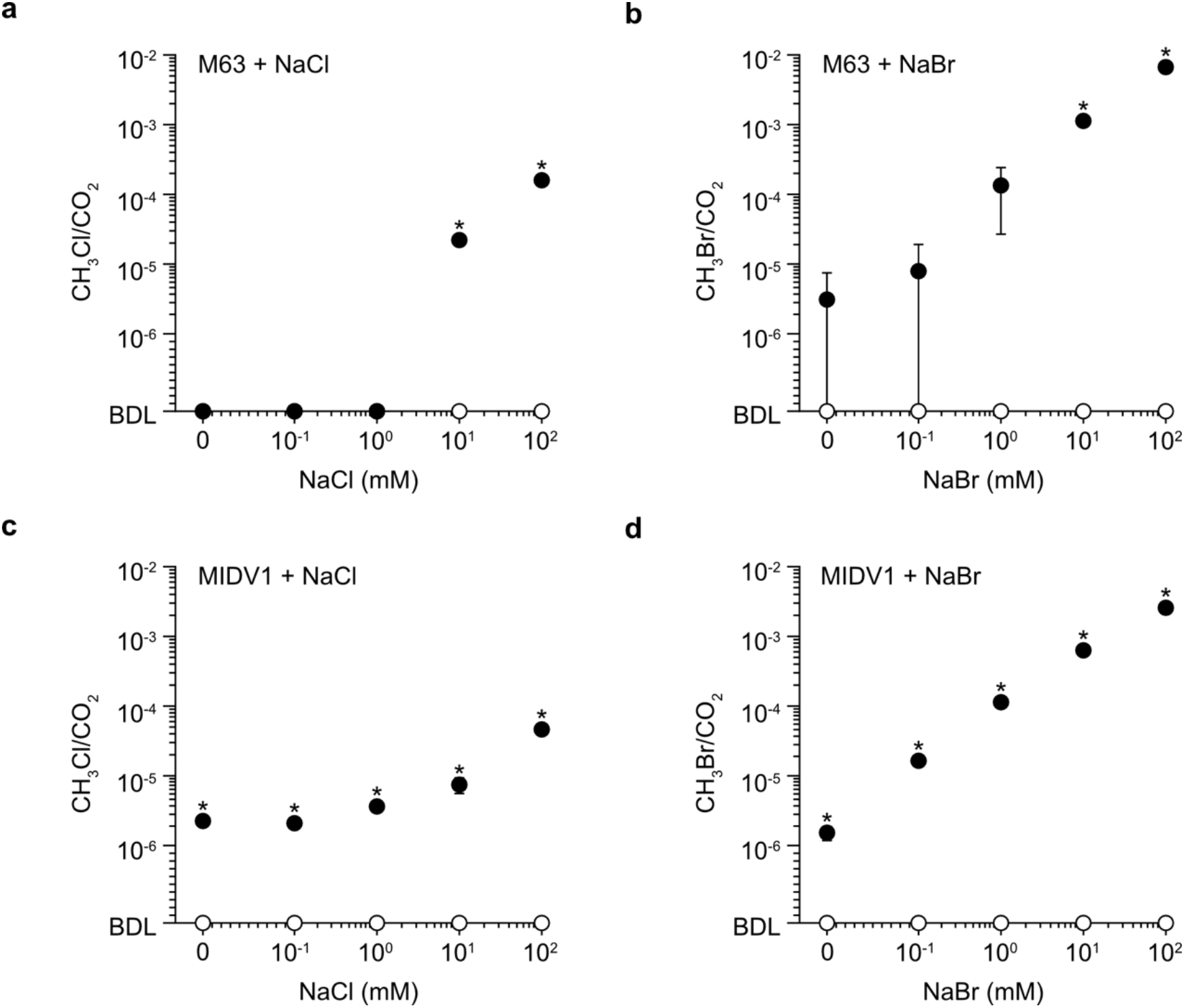
Effect of halide augmentation on the indicator gas signal in soil. **(a)** CH_3_Cl and **(b)** CH_3_Br signals in the headspace of autoclaved B horizon soil (1 gram) hydrated to 32% field capacity with M63 containing 10^7^ CFU/gram soil of MG1655-*mht* (●) or wildtype MG1655 (○) and varying concentrations of halides. The CH_3_X signal was measured after a 24-hour incubation at 22°C and normalized to CO_2_ to account for cellular respiration. Only samples supplemented with ≥10 mM halide presented a CH_3_X signal significantly above background (*, p < 0.01). **(c)** CH_3_Cl and **(d)** CH_3_Br signal from autoclaved B horizon soil (1 gram) hydrated to 64% field capacity with MIDV1 containing 10^7^ CFU/gram soil of MG1655-*mht* (●) or wildtype MG1655 (○) and varying concentrations of halide (0 to 100 mM). The CH_3_X signal was significantly above background at all halide concentrations (*, p < 0.01). Data represent the average of three biological replicates, while error bars represent ±1 standard deviation. p-values were calculated using a two-sided, independent *t* test. BDL, below detection limit.

Soils hydrated with minimal growth media, such as M63, exhibit hydrologic properties very different from those in natural soils because the osmotic pressure of minimal growth medium is higher than the aqueous phase of soils *in situ*. One way to overcome this challenge is to hydrate soils with diluted minimal medium, such as MIDV1^16^. To determine how hydration at a more realistic osmotic pressure affects the output from gas-reporting microbes, we evaluated the signal from soils hydrated to 64% field capacity using MIDV1 containing MG1655-*mht* (10^7^ CFU/gram soil). In the absence of NaCl or NaBr supplementation (Figures 1c-d), we found that indicator gas signals were significantly above the CH_3_X background (p < 0.01, two-sided *t* test). For both halides, the headspace gas signal increased with the concentration of halides added to soil. A comparison of gas signals from soil hydrated with M63 and MIDV1 medium revealed that the CH_3_Br signal is higher than the CH_3_Cl signal when similar levels of each halide salt are added. This trend is interpreted as arising because the *Batis maritima* MHT used for this study has a lower K_m_ for bromide (18.5 mM) compared to chloride (155 mM)^30^.

In the environment, soil microbes live under conditions that are frequently nutrient-limiting to model synthetic biology chassis, such as *E. coli*^31^. To better understand how growth limitations affect gas production, we incubated MG1655-*mht* in soil hydrated to 64% field capacity with low-osmolarity MIDV1 (20 mM NaBr) containing or lacking nitrogen (Figure 2a). Under nitrogen-deficient (-N) conditions, we found that the maximum gas signal was 44% of that observed in samples containing nitrogen following a 24-hour incubation. To evaluate how cell titer and supplemental halides affected gas signal under nitrogen-limiting conditions, we quantified the signal from soil containing different titers of MG1655-*mht* that had been hydrated to the same water content with MIDV1 (-N) containing a range of NaBr concentrations (Figures 2b, S3). The CH_3_Br signal correlated with both cell titer and supplemental halide concentration following a 24-hour incubation. Across the range of conditions tested, both NaBr concentration and cell titer had similar effects on the CH_3_Br signal, such that an order of magnitude increase in either the cell population or NaBr concentration increased the gas signal by similar amounts.

**Figure 2.**
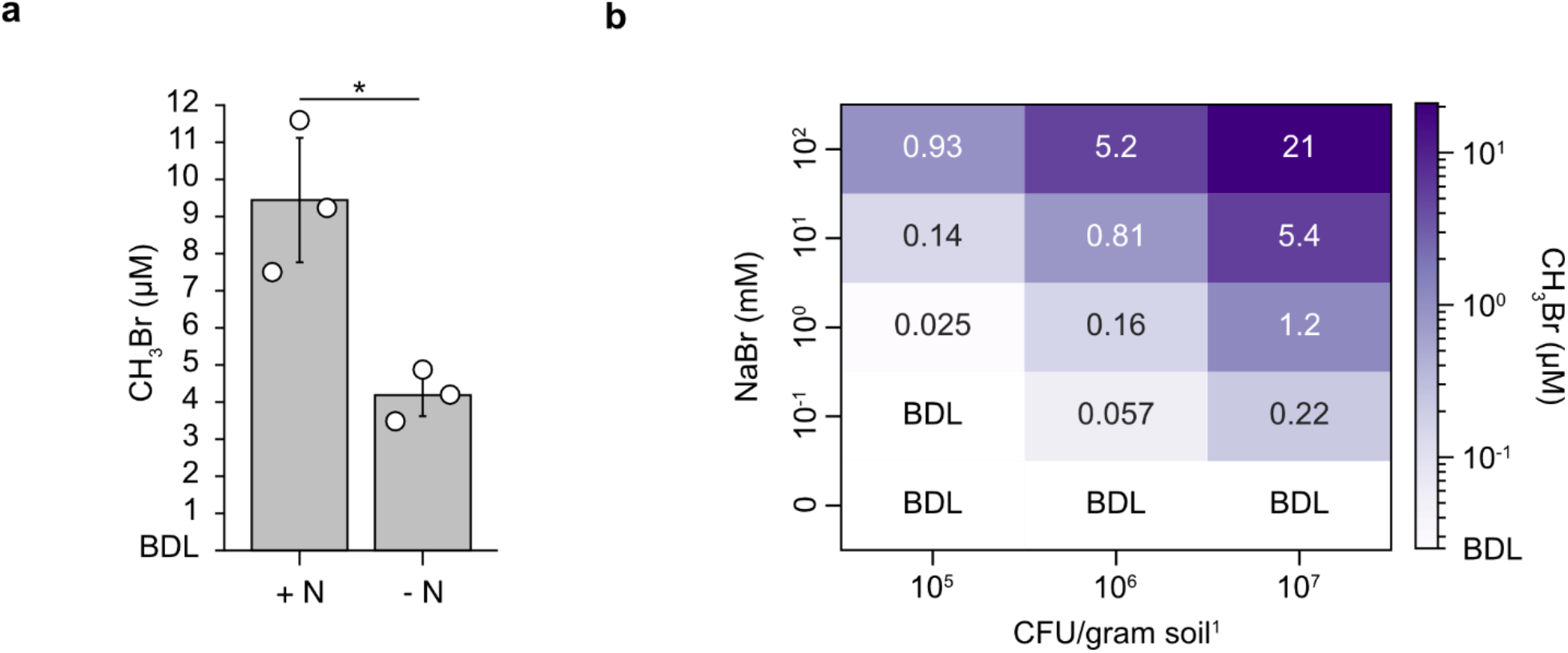
Nitrogen limitation decreases the gas reporter signal in soil. **(a)** CH_3_Br signal presented by MG1655-*mht* (10^7^ CFU/gram soil) in soil containing (+N) or lacking (-N) nitrogen. Autoclaved B horizon soil (1 gram) hydrated to 64% field capacity with MIDV1 (20 mM NaBr) was incubated at 22°C for 24 hours before headspace gas analysis. Indicator gas signals from samples lacking nitrogen were significantly decreased from samples containing nitrogen (p = 0.01). **(b)** CH_3_Br signal presented by varying titers of MG1655-*mht* in autoclaved B horizon soil (1 gram) hydrated to the same field capacity with MIDV1 (-N) containing varying amounts of NaBr. Samples were incubated at 22°C for 24 hours prior to gas analysis. Values shown represent the average of three biological replicates, with the individual data points are shown as open circles. p-values were calculated using a two-sided, independent *t* test. Error bars represent ±1 standard deviation. BDL, below detection limit.

### Gas signal dynamics in soil

Many soil processes are associated with hot spots and hot moments that occur sporadically in the environment^1,2^ and may fluctuate over timescales (days to seasons) that exceed short incubations (hours to days) typically used for gas biosensors^8,12^. As such, it is important to understand indicator gas persistence over longer timescales. To better understand gas signal persistence, we measured headspace gas accumulation in liquid and soil samples over a one-week incubation.

We first evaluated the CH_3_Br signal persistence in liquid cultures containing MG1655 *E. coli* that constitutively express MHT. We incubated liquid cultures (10^6^ CFU/mL) in M63 medium (-N; 100 mM NaBr) for one week at 37°C in a shaking incubator and measured the headspace CH_3_Br that accumulated following different incubation durations (Figure 3a). We also measured CO_2_ accumulation so that we could compare the indicator gas signal dynamics with respiration and metabolic activity (Figure 3b). In liquid culture, the peak CH_3_Br signal was observed following 1 day of incubation, while the peak CO_2_ signal was observed after 2 days. The CH_3_Br signal declined following the peak, while the CO_2_ signal remained relatively constant at later time points. The number of viable cells decreased by approximately three orders of magnitude following one week of incubation (Figure S4). To better understand the mechanism underlying the CH_3_Br decline, we added a CH_3_Br standard to liquid cultures containing or lacking wildtype *E. coli* (Figure S5a). With both measurements, we found that CH_3_Br decreased exponentially at similar rates. These findings suggest that the decline is driven by chemical (OH^-^ and Cl^-^) substitution of CH_3_Br in the liquid phase^32–34^. No significant difference in cell viability was observed between *E. coli* cultures containing or lacking CH_3_Br (Figure S5b). This latter finding suggests that the amount of CH_3_Br produced by *E. coli* expressing MHT does not affect cell fitness.

**Figure 3.**
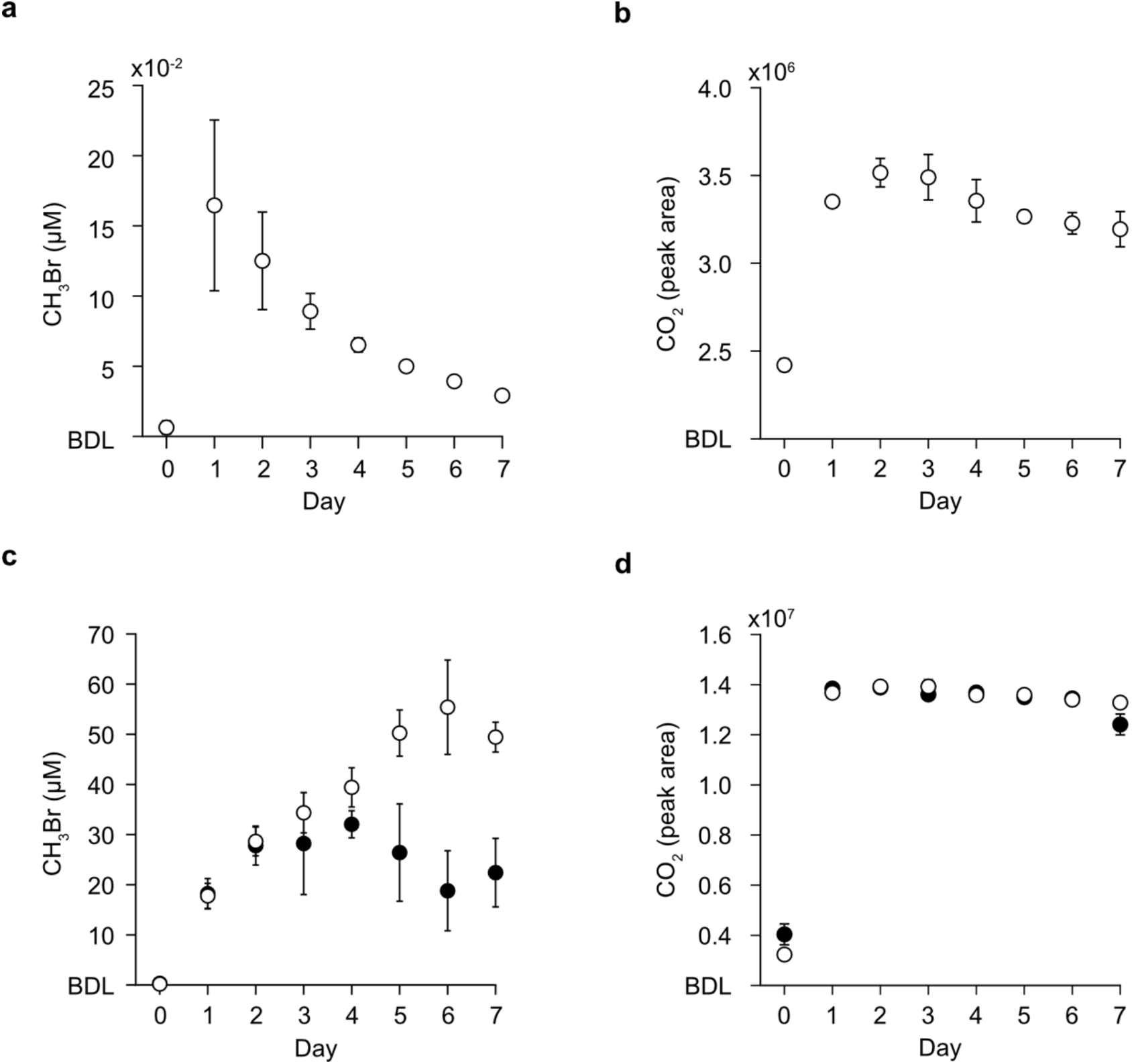
Indicator gas signal persistence. **(a)** CH_3_Br and **(b)** CO_2_ accumulation in liquid samples containing MG1655-*mht* (10^6^ CFU/mL) in M63 (1 mL; -N, 100 mM NaBr). Identical samples were incubated in sealed GC-MS vials at 37°C while shaking at 250 rpm, and matched samples were sacrificed for gas analysis every 24 hours. **(c)** CH_3_Br and **(d)** CO_2_ generated by MG1655-*mht* (10^7^ CFU/gram soil) added to either unsterilized (●) or autoclaved (○) B horizon soil (1 gram) hydrated to 64% field capacity with MIDV1 (-N; 20 mM NaBr). Samples were incubated without shaking at 22°C in sealed vials. Data represent the average of three biological replicates. Error bars represent ±1 standard deviation. BDL, below detection limit.

To evaluate whether CH_3_Br persistence is also affected by soil, we evaluated the dynamics of a CH_3_Br signal generated by MG1655-*mht* (10^7^ CFU/gram soil) in B horizon soil hydrated to 64% field capacity with MIDV1 medium (-N; 20 mM NaBr). We performed these experiments in soil containing a native soil microbiome and soil subjected to two sequential autoclaving cycles. For these measurements, we monitored total CH_3_Br and CO_2_ accumulation at 22°C each day for a week (Figures 3c-d). The indicator gas signals observed in the autoclaved and untreated soils were similar until day 4. After that time, the matrices containing a soil microbiome presented a ∼2-fold lower CH_3_Br signal, which was not significantly different from the twice-autoclaved soils (p > 0.05, two-sided *t* test). In both samples, the CO_2_ signal increased after the first day and remained stable thereafter. Taken together, these findings show that indicator gas signals are degraded in environmental matrices containing a microbiome.

To better understand how changes in gas production relate to microbial growth and fitness, we extracted and counted cells from the unsterilized soil at each time point. The number of CFU on agar plates containing an antibiotic selection for MG1655-*mht* peaked after one day and remained consistent over time (Figure S6). These results suggest that the observed CH_3_Br decline in living soils is primarily driven by increased degradation of the CH_3_Br signal rather than decreased viability of *E. coli*. In addition to abiotic degradation, microbial consumption of CH_3_X by methylotrophs has been documented in a wide range of soils as a CH_3_X sink in the environment^29,35–37^.

### Effect of sample concentration on the limit of detection

We next sought to quantify how background CH_3_Br production from unsterilized soils was influenced by supplemental halides. To achieve the analytical sensitivity necessary to detect ambient levels of CH_3_Br, we implemented a TD system that concentrates headspace gas prior to GC-MS analysis. We performed incubations in 2.8 L flasks, sampled air from the flask headspace into evacuated 0.5 L sample canisters, pumped the gas sample from the canisters to a TD system with a sorbent-packed cold trap that collects CH_3_X, and analyzed the CH_3_X signal following heat-induced desorption of the trap onto a GC-MS (Figure S7). We compared standard curves generated by TD-GC-MS and GC-MS alone and found that indicator gas concentration using TD increased our sensitivity of detecting CH_3_Br and CH_3_Cl by more than two orders of magnitude (Figures S8a-b). To understand ambient CH_3_X variability in the lab where we performed incubations, we measured CH_3_Br and CH_3_Cl over six months. From June to December 2021, we observed an average of 954 ppt CH_3_Cl (±147 ppt, mol/mol) and 26 ppt CH_3_Br (±8 ppt, mol/mol) in the lab (Figure S8c-d). These laboratory values are 1.7- to 3.7-fold higher than mole fractions observed in the environment, ∼550 ppt CH_3_Cl and ∼7 ppt CH_3_Br^26^, suggesting that the methyl halide background will be lower in natural environments.

To determine whether soil hydration increased CH_3_Br levels above the ambient background, we compared CH_3_Br in the headspace of flasks containing either unsterilized or twice-autoclaved soil from the B horizon of the Alfisol. We hydrated the soil to 64% field capacity with MIDV1 medium (-N; 100 mM NaBr) and incubated the soil in sealed flasks for 48 hours at 22°C. This incubation time was chosen because it yielded the maximum signal in week-long incubations. Both soil samples presented a significant CH_3_Br signal (p < 0.01, two-sided *t* test) compared to ambient levels (Figure 4a). Furthermore, over this short time scale, the living soil accumulated significantly more CH_3_Br than the autoclaved soils. Other studies have observed that sterilization treatments significantly alter net CH_3_X flux from soils, affirming that biological processes dominate CH_3_X fluxes *in situ*^29,38–40^.

**Figure 4.**
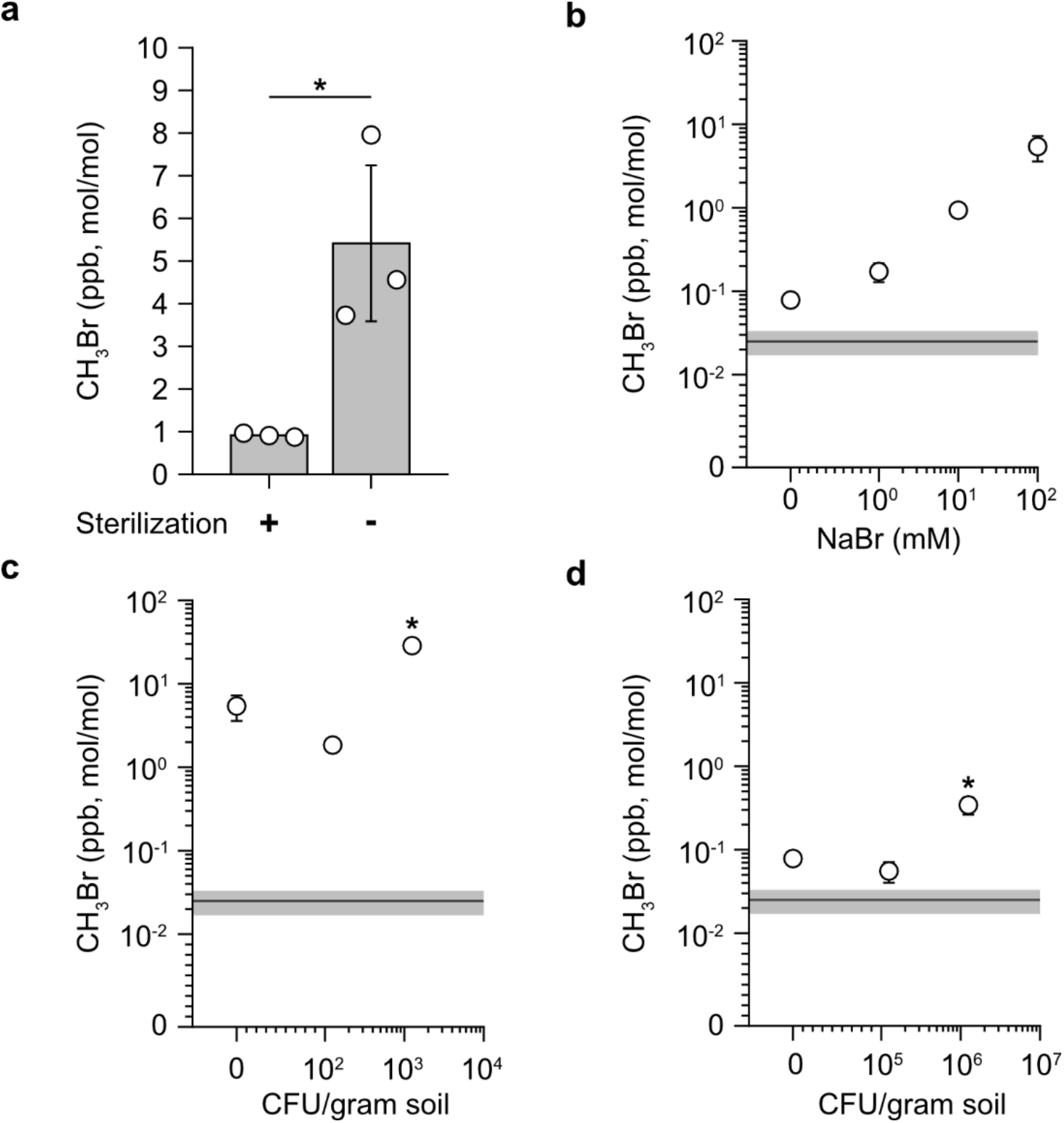
Effect of gas concentration on signal detection. **(a)** CH_3_Br from an autoclaved (+) and unsterilized (-) B horizon soil (1 kg) that had been hydrated to 64% field capacity with MIDV1 (-N; 100 mM NaBr) and incubated for 48 hours at 22°C. Unsterilized soil presents a significantly higher signal (*, p = 0.025) than autoclaved soil. **(b)** Effect of NaBr augmentation on CH_3_Br produced by unsterilized B horizon soil that was similarly hydrated. All samples presented significantly higher CH_3_Br (p < 0.01) than ambient, which is shown as a line. **(c)** Effect of MG1655-*mht* titer on the gas signal from unsterilized B horizon soil hydrated with MIDV1 (-N, 100 mM NaBr) to the same water content. A MG1655-*mht* titer of 10^3^ CFU/gram soil was required to present a signal significantly above the no-cell background (*, p = 0.001). **(d)** Effect of MG1655-*mht* titer on the headspace gas signal from unsterilized B horizon soil hydrated to the same water content with MIDV1 (-N) lacking supplemental halides. A higher cell titer (10^6^ CFU/gram soil) was required for a significant signal above the no-cell control (*, p = 0.01). Data represent the average of three biological replicates, while error bars represent ±1 standard deviation. Average ambient concentration (dark gray line) is plotted ±1 standard deviation (light gray bar). p-values were calculated using a two-sided, independent *t* test.

We next sought to determine the effect of NaBr supplementation on soil production of CH_3_Br, since the Alfisol being used for microbial incubations contains naturally low bromide levels. To do this, we incubated unsterilized B horizon soil hydrated to 64% field capacity with MIDV1 medium (-N) that contained a range of NaBr concentrations. After 48 hours at 22°C in the absence of a gas-reporting microbe, all samples presented significantly higher CH_3_Br levels than the ambient background (p < 0.01, two-sided *t* test) (Figure 4b). CH_3_Br accumulation also increased with increasing halide supplementation, indicating that halide availability is a limiting factor for background CH_3_Br production in these soils.

To determine how many gas-producing cells are required to present a signal above the soil background, we hydrated the unsterilized B horizon soil to 64% field capacity with MIDV1 (-N) containing or lacking NaBr (Figure 4c-d). When supplemented with 20 mM NaBr, 10^3^ CFU/gram soil of MG1655-*mht* was sufficient to produce a signal that was higher than the soil background (p = 0.001, two-sided *t* test). In the absence of halide supplementation, 10^6^ CFU/gram soil was required to produce a significant signal (p = 0.01, two-sided *t* test). Taken together, these results show that halide concentration affects the magnitude of both the background CH_3_X produced by the soil and the signal produced by engineered cells added to the soil.

### Sensing a sugar in living soil

Our results with microbes that constitutively produce an indicator gas suggested that biosensing cells may be able to report on chemicals when present at similar low titers in a living soil. To test this idea, we evaluated the gas signal from MG1655 cells containing an MHT gene whose expression is regulated by the lac repressor (MG1655-IPTG)^41^. In this strain, MHT expression and CH_3_X biosynthesis is induced by isopropyl ß-D-1-thiogalactopyranoside (IPTG). To first investigate whether soil affects IPTG bioavailability, we added MG1655-IPTG (10^8^ CFU/gram soil) to GC vials containing or lacking unsterilized B horizon soil that was hydrated to 64% field capacity with MIDV1 (-N; 20 mM NaBr) containing varying IPTG concentrations. We incubated the samples at 22°C for 24 hours before analyzing the headspace gas (Figure 5a). The dynamic range of IPTG sensitivity was similar in liquid and soil incubations, demonstrating that IPTG exhibits high bioavailability in the soil. To benchmark the gas output from the IPTG biosensor to MG1655-*mht*, we compared the CH_3_Br signal from soil samples containing similar titers (10^8^ CFU/gram soil) of each strain. We performed these experiments in unsterilized B horizon soil hydrated to 64% field capacity with MIDV1 (-N; 100 mM NaBr) containing or lacking IPTG. After a 24-hour incubation at 22°C, MG1655-IPTG produced ∼2-fold more CH_3_Br in the presence of IPTG compared to MG1655-*mht* (Figure 5b). MG1655-IPTG also presented a CH_3_Br signal in the absence of IPTG, although IPTG addition increased the signal by 116-fold in the soil headspace.

**Figure 5.**
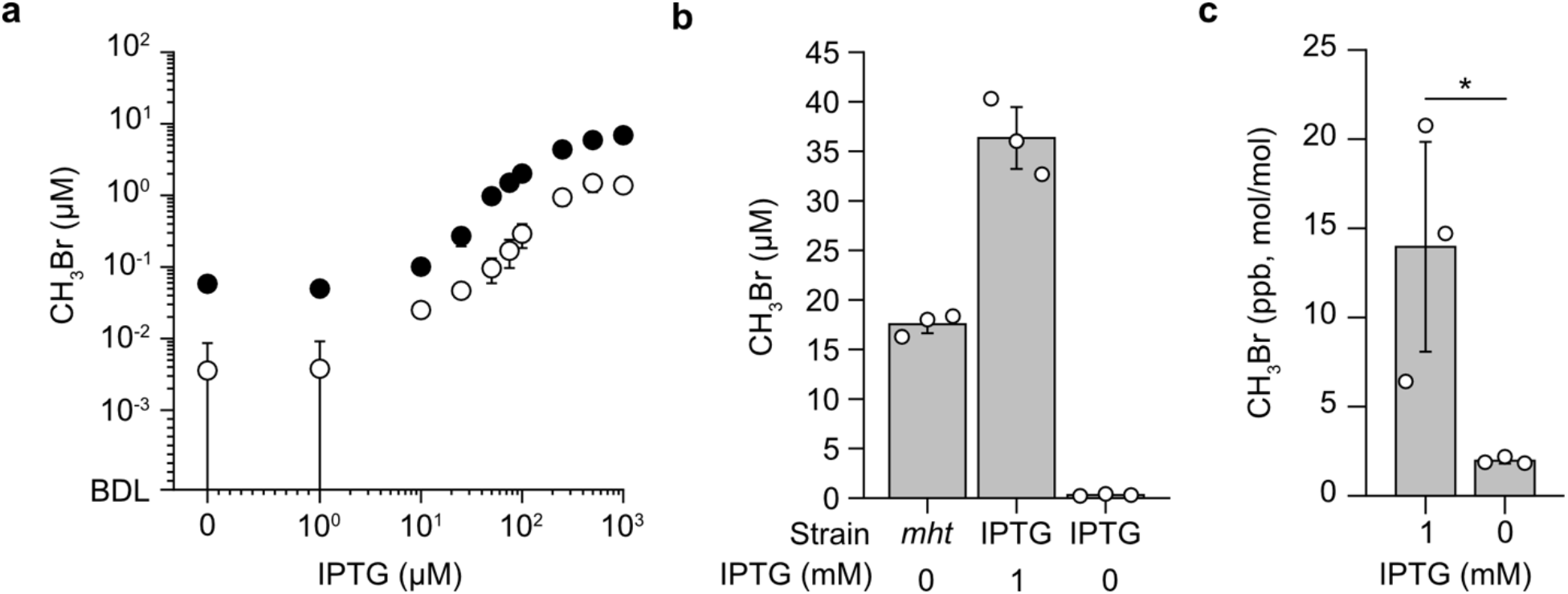
Chemical sensing in soil. **(a)** Effect of IPTG on CH_3_Br produced by MG1655-IPTG. MIDV1 (-N; 20 mM NaBr) with chloramphenicol (34 µg/mL) and varying IPTG was added to 2 mL GC-MS vials containing (●) or lacking (○) unsterilized B horizon soil (1 gram). Cells (10^8^ CFU) in the same medium were added to each vial, yielding soil samples hydrated to 64% field capacity. Headspace gas was measured after 24 hours incubation at 22°C. **(b)** Indicator gas signals from MG1655-*mht* (*mht*) and MG1655-IPTG (IPTG) at a titer of 6×10^7^ CFU in unsterilized B horizon soil (1 gram) hydrated to the same water content with MIDV1 (-N; 100 mM NaBr) containing or lacking IPTG. Headspace gas was measured following 24 hours of incubation at 22°C **(c)** TD enables detection of IPTG in an unsterilized B horizon soil (1 kg) hydrated to 64% field capacity with MIDV1 (-N; 100 mM NaBr) and containing 10^3^ CFU/gram soil MG1655-IPTG. Headspace gas was measured using TD-GC-MS following a 48-hour incubation at 22°C. Samples with IPTG produced significantly more CH_3_Br than samples lacking IPTG (*, p = 0.045). Data represent the average of three biological replicates. p-values were calculated using a two-sided, independent *t* test.

To investigate if the IPTG sensor could produce a detectable signal in a living soil when present at the lowest titer where MG1655-*mht* presented a signal, we added MG1655-IPTG at a titer of 10^3^ CFU/gram soil and hydrated untreated B horizon soil to 64% field capacity with MIDV1 (-N; 100 mM NaBr) containing or lacking IPTG (Figure 5c). After 48 hours of incubation at 22°C, samples containing IPTG presented a 7-fold higher CH_3_Br signal than samples lacking IPTG, and the signal with IPTG was significantly higher than without IPTG (p = 0.045, two-sided *t* test). CH_3_Br production in the absence of IPTG was not significantly different from background soil production (p = 0.83, two-sided *t* test).

### Implications

Our results demonstrate that non-invasive biosensing with minimal perturbation of the environmental matrix can be achieved with cell titers as low as 10^3^ cells/gram soil. Additionally, biosensing at this titer was accomplished in a soil containing an environmental microbiome. The net indicator gas signal presented by a microbe expressing an MHT depends upon many parameters including engineered cell titer, soil halide concentration, and nutrient availability. While a gas signal from biosensing cells could be obtained from a living Alfisol without adding halides, the number of cells required to observe a signal above laboratory background was sensitive to halide supplementation. The cell titers required in the presence and absence of halide supplementation, 10^3^ and 10^6^ cells/gram soil respectively, are both smaller than the titer of cells (10^7^ to 10^12^ cells/gram soil) observed in soil microbiomes^1,21,22^. As the soil analyzed in this study has low halide concentrations, soils with higher halides may require fewer biosensors to yield a signal in the absence of halide supplementation. The low limit of detection achieved using TD-GC-MS suggests that indicator gas reporting could be used to monitor the gene expression of native and engineered cells present in hard-to-image materials, ranging from microbiome-containing environmental matrices like soils, sediments, and wastewater to synthetic living materials.

The limit of detection for gas-reporting biosensors is expected to vary across materials, and future applications of these biosensors should always include control measurements to establish the background methyl halide levels in each matrix. In our Alfisol incubations, hydrated soils were net sources for CH_3_Br following a two-day incubation. This gas production is interpreted as primarily arising from fungal methyltransferase activity induced by hydrating the soil with medium containing unnaturally high concentrations of halides^29,42,43^. While this background can be minimized by decreasing the halide supplementation, decreasing the halides also attenuates the signal produced by the engineered microbes. The application of gas biosensors in other materials that present background methyl halides, including marine^44,45^ and lacustrine^46,47^ sediments and samples containing vegetation^48,49^, will also require control measurements to account for natural CH_3_X fluxes.

Our results also demonstrate that the limit of detection for gas biosensors depends upon the incubation duration. This variation arises because methyl halide production and consumption processes vary dynamically in the matrix. When soil was inoculated with engineered cells and incubated for a week, the CH_3_X signal showed a greater decline in unsterilized soils compared to autoclaved soils. We interpret this trend as arising from bacterial degradation, which has been suggested as a major sink for methyl halides in soils^29,35–37^. Abiotic degradation processes including hydrolysis and nucleophilic substitution can also decrease indicator gas signals^32–34^, the latter of which may be particularly important when gas biosensors are deployed in high salinity environments. Abiotic methyl halide production in soils with high organic matter content has also been observed^29,50^, indicating that both microbiome and matrix effects are important for determining net methyl halide accumulation over time. Further studies will be needed to elucidate how environmental parameters influence indicator gas signals across different environmental materials, microbiomes, and time scales.

Indicator gas biosensor outputs are expected to be matrix- and treatment-dependent. As such, the optimal supplementation and hydration to maximize the signal-to-noise ratio for gas biosensors will vary in each environmental matrix. The soils used in this study have halide concentrations that are more than one order of magnitude below the estimated global averages^17^, suggesting that other matrices may not require supplemental halides for high indicator gas production. Soils with naturally elevated halides that experience moisture regimes favorable to microbial activity are expected to present robust indicator gas production when gas biosensors are introduced into those soils. For example, coastal agricultural soils often have high salinity because they experience rainfall of marine origin with a high salt aerosol content and because seawater can intrude into the ground water^51^. Water content and soil texture will also affect indicator gas signal by impacting biosensor viability and gas diffusion. Under extreme dry conditions, microbial metabolic activity is expected to be attenuated, thereby decreasing indicator gas production rates, while water saturated conditions are expected to hamper gas diffusion from soil pores to the headspace^52^. Natural CH_3_X fluxes in soils are also known to vary with temperature^40,53^, hydration^48,54^, organic matter content^29,50^, halide concentration^17^, redox properties^39,54^ and microbiome^29,40^. Additional studies are needed to understand how these processes affect total indicator gas accumulation in the headspace of different matrices.

In future studies, several approaches could be used to improve the signal-to-noise ratio when applying indicator gas reporters in low-halide environmental materials like the Alfisol used here. MHT mutants with greater affinity for halide ions could be developed to improve gas reporting in low-halide environments by increasing the signal from engineered cells under conditions that minimize background. Selections have been reported for methyl transferases that could be applied to identify MHTs with improved substrate affinities^55^. Deep sequencing a library of MHT mutants before and after selection^56^ could identify mutations that improve cellular fitness in low halide growth conditions. Additionally, gas biosensor signals could be better differentiated from environmental background by combining gas concentration with ^13^C labelling of biosensor cells and using isotope ratio mass spectrometry to enable detection of trace methyl halide isotopologue abundances^29^. Finally, genetic circuits encoding gas reporters could be transferred into native cellular chassis to reduce the impact of non-native inoculants on soil microbiome structure and decrease the need for supplemental nutrients that may alter matrix processes^23–25^. We expect that these advancements, combined with the ultrasensitive detection described here, will facilitate gas-reporting biosensing in a wide variety of environmental and synthetic matrices.

## METHODS

### Materials

Chloramphenicol was from MilliporeSigma, liquid methyl halide standards were from Restek, and gas-phase methyl halide standards were from Airgas. IPTG was from Research Products International, sodium halides used to create ion chromatography standard curves were from MilliporeSigma, and all other chemicals were from VWR, MilliporeSigma, Fisher, Restek, Apex Biosciences, Research Products International, or BD Biosciences. Enzymes and reagents for molecular biology were from New England Biolabs and Qiagen. Gas tight vials (2 mL) were from Phenomenex, and gas sample canisters were from LabCommerce.

### Strains and plasmids

Table S3 lists the plasmids and strains used in this study. MHT was constitutively expressed in MG1655 using a previously described strain (MG1655-*mht*) that expresses *Batis maritima* MHT from the chromosome^8^ or MG1655 transformed with a plasmid (pEMF051) that uses a strong constitutive promoter^57^ to express a fusion of sfGFP^58^ and MHT^8^ genes linked via (GGGGS)_2_. MHT was conditionally expressed in MG1655 transformed with a plasmid (pLC7) that regulates expression of MHT using a promoter regulated by the lac repressor (LacI)^41^. All plasmids were generated using Golden Gate cloning^59^ and sequence verified. pLC7 and pEMF051 contain a chloramphenicol resistance marker and the ColE1 origin. All molecular biology was performed using *E. coli* XL1-Blue.

### Microbial growth

For cloning and strain maintenance, cells were grown in Lysogeny Broth (LB) at 37°C. Studies analyzing CH_3_X production in liquid culture used a modified M63 medium^8^ (pH = 7.0) containing 1 mM magnesium sulfate, 0.2% w/w glucose, 0.00005% w/w thiamine, 0.05% w/w casamino acids, 20% w/w M63 salt stock, and the indicated concentrations of NaCl or NaBr. Experiments in low-osmolarity medium used modified MIDV1 medium^16^ (pH = 7.0), which contained 1 mM magnesium sulfate, 0.2% w/w glucose, 0.00005% w/w thiamine, 0.05% w/w casamino acids, 0.0125% w/w M63 salt stock, and the indicated concentrations of NaCl or NaBr. The M63 salt stock contained 75 mM ammonium sulfate, 0.5 M monobasic potassium phosphate, and 10 µM ferrous sulfate^12^. N-deficient media lacked ammonium sulfate and casamino acids. The optical density of cultures at 600 nm (OD_600_) was monitored using a spectrophotometer (DS-11, DeNovix). To calibrate OD_600_ with cell titer, MG1655 cultures of varying OD_600_ were incubated overnight on LB-agar plates, and the resulting colonies were counted.

### GC-MS analysis

For headspace gas analysis without sample concentration, samples (soil and liquid culture) were added to 2 mL gas-tight vials (AR0-37L0-13, Phenomenex) and crimped with VEREX tops (AR0-5710-13, Phenomenex). After incubation, headspace gas was quantified using an Agilent 7820 GC connected to an Agilent 5977E MS or an Agilent 8890 GC connected to an Agilent 5977B MS, both with autosamplers (7693A, Agilent). Both systems contained a DB-VRX capillary column (20 m, 0.18 mm ID, 1 µm film; Agilent) and a 100 µL gastight syringe (G4513-80222, Agilent) for sampling. The Agilent 7820 GC used a 50:1 split ratio for headspace gas injections (50 µL), a column flow of 0.9 mL, and an oven program that held at 45°C for 1.3 min. The Agilent 8890 GC used a 55:1 split ratio for headspace gas injection (50 µL), a column flow of 1.1 mL/min, and an oven program that held at 45°C for 84 s before ramping to 60°C at 36°C/min and then holding at 60°C for 9 s. In both systems, MS analysis was performed using selected ion monitoring (SIM) for CH_3_Br (*m/z* = 94 and 96), CO_2_ (*m/z* = 44 and 45), and CH_3_Cl (*m/z* = 50 and 52). Agilent MassHunter WorkStation Quantitative Analysis software was used to quantify the peak area of the major isotopologue of each chemical, while the minor isotopologue was used as a qualifier. For experiments performed in 2 mL vials, CH_3_X standard curves were generated at each experimental condition by serially diluting 2 mg/mL liquid CH_3_Br (Restek, 30253) or CH_3_Cl (Restek, 30267) into growth medium. Standards were added to GC vials containing (1 gram) or lacking soil and allowed to equilibrate under experimental conditions for at least 4 hours before measurement.

### Soil sampling

Soil was collected from property owned by Nature’s Way Resources, a compost facility located in Conroe, TX on February 17, 2020. Soil was sampled from a part of the property not used for compost production, undisturbed since at least 1984 (John Ferguson, pers. comm.). Samples were acquired from the A (0-10 cm) and B (10-58 cm) horizons separately. Each horizon was thoroughly mixed on site, transported to the lab, and immediately stored at -20°C before processing. Samples were oven-dried at 60°C, passed through a 2 mm sieve to remove roots and large fragments, and placed in 12-gallon plastic storage bins (Home Depot) prior to storage at room temperature, dry, to minimize organic matter degradation. To sterilize soils, samples were autoclaved for 45 minutes at 120°C, allowed to rest overnight, and autoclaved again.

### Soil characterization

Soil was ground with an agate mortar and pestle prior to chemical analysis. Total C, H and N were measured by catalytic combustion using an elemental analyzer (ECS 4010, Costech) and subsequent chromatographic separation and detection of CO_2_, H_2_O, and N_2_. To quantify organic C, inorganic C was removed using 7M HCl acid fumigation in open-top silver capsules^60^. Water extractable halides were measured using an ion chromatograph (Dione ICS-2100, Thermo Scientific). Soil (10 g) was suspended in deionized water (50 mL) and mixed on an orbital shaker (OS-500, VWR) for 2 hours. The suspension was centrifuged, and the supernatant was filtered through a 0.22 µm syringe filter. Standard curves for the halides were generated using solutions containing 1, 10, or 100 mg/L mixtures of NaCl, NaBr, and NaI. Soil pH was measured using a 1:2 soil/water mixture (w/w) after 30 min equilibrium with atmospheric CO_2_.

### Effect of halides on gas production in soil

MG1655-*mht* and MG1655 were grown to mid-log phase in M63 medium lacking halides. Cells were resuspended in M63 (100 µL) supplemented with a range of NaBr or NaCl (0 to 100 mM) and added to 2 mL GC vials containing twice-autoclaved B horizon soil (1 gram) to a water content representing 32% field capacity; the final cell density was 3×10^7^ CFU/gram soil. For samples hydrated with MIDV1, cultures were grown to mid-log phase in M63 lacking halides and washed thrice in MIDV1 lacking halides. Cells were resuspended in MIDV1 (200 µL) supplemented with varying NaBr or NaCl (0 to 100 mM) and added to GC vials containing twice-autoclaved B horizon soil (1 gram) to a water content of 64% field capacity; the final cell titer was 6×10^7^ CFU/gram soil. Samples were sealed and incubated at 22°C for 24 hours before headspace gas analysis by GC-MS.

### Effect of N limitation on the gas signal

MG1655-*mht* were grown to mid-log phase in M63 lacking halides, washed thrice, and resuspended in MIDV1 (20 mM NaBr) containing or lacking nitrogen (-N). Growth medium (200 µL) containing 6×10^7^ CFU was added to GC vials containing twice-autoclaved B horizon soil (1 gram) for a water content of 64% field capacity. For samples containing varying CFU and NaBr, cultures of MG1655-*mht* were grown to mid-log phase and washed thrice in MIDV1 lacking nitrogen and halides. Cells were resuspended in MIDV1 (-N, 0 to 100 mM NaBr) and added to twice-autoclaved B horizon soil (1 gram) to a water content representing 64% field capacity and cell titer of 6×10^5^, 6×10^6^, or 6×10^7^ CFU/gram soil. Samples were sealed and incubated for 24 hours at 22°C before headspace gas analysis by GC-MS.

### Signal persistence measurements

To assess CH_3_Br stability in liquid cultures, MG1655 carrying a plasmid for constitutive expression of the sfGFP-MHT fusion (pEMF051) were grown to mid-log phase in M63 (100 mM NaBr) containing chloramphenicol (34 µg/mL). Cells were washed thrice, resuspended in M63 (1 mL; -N, 100 mM NaBr) containing chloramphenicol (34 µg/mL), and added to 2 mL GC vials at a cell density of 10^6^ CFU/mL. Samples were capped and incubated at 37°C while shaking at 250 rpm. Each day, matched samples were sacrificed for headspace gas analysis. Immediately after sampling, serial dilutions of each sample were plated on LB-agar medium containing chloramphenicol (34 µg/mL) and incubated overnight at 37°C to enable CFU counting. To assess soil effects on CH_3_Br signal, MG1655-*mht* was grown to mid-log phase in M63 lacking halides, washed thrice, and resuspended in MIDV1 (-N, 20 mM NaBr). Resuspended cells (200 µL) were added to twice-autoclaved or unsterilized B horizon soil (1 gram) at a final titer of 6×10^7^ CFU/gram soil and a water content of 64% field capacity. Samples were sealed and incubated statically at 22°C. Each day, matched samples were sacrificed for headspace gas analysis.

### Incubation chambers

Fernbach flasks (4420, Pyrex) were fitted with two-holed rubber stoppers (size 13; 14-140Q, Fisher) and sealed around the rim using silicone tape (VP1014, Tommy Tape). A sampling tube assembly was fit into one stopper hole, which consisted of a 2-inch segment of ¼’’ Sulfinert-treated stainless steel tubing (RE22902, Restek), a 2 µm Siltek-coated frit filter (24171, Restek), a plug valve (SS-4P4T, Swagelok), and a second segment of the same tubing. A second tube assembly lacking the frit filter was fitted into the other hole to enable pressure re-equilibration after sampling. To minimize CH_3_X and microbiological carryover between samples, flasks and stoppers were autoclaved, sampling tubes were soaked in 70% ethanol, and stoppers were degassed 3x in a vacuum chamber refilled with N_2_ (Airgas). Samples for TD-GC-MS analysis were collected from the chambers using 0.5 L gas sampling canisters (X20.5L, LabCommerce) that had been evacuated to <10^−4^ mbar using a dry scroll pump (nXDS10i, Edwards). During sampling, canisters were allowed to equilibrate with the chamber headspace for ∼15 seconds. To minimize analyte carryover, canisters were purged with 10 psig He (HE UHP300, Airgas) between experiments.

### Thermal desorption

Canisters were connected to an autosampler (CIA Advantage-x, Markes) using transfer lines heated to 120°C. Samples of varying volume (10 to 400 mL) were transferred from the 0.5 L gas canisters to the TD system at 50 mL/min and passed through a water condenser (Kori-xr, Markes) with a cold trap (−30°C) to remove water. The TD (UNITY-xr, Markes) was equipped with a PAMS focusing trap (U-T20PAM-2S, Markes) that was purged for 1 min with 50 mL/min N_2_ before sample loading at 20°C. Finally, the trap was desorbed for 1 min with a split flow of 2 mL/min at 280°C by ultrapure He carrier gas onto an Agilent 7890B GC fitted with a PoraPLOT Q capillary column (25 m, 0.25 mm, 8 µM film) and connected to an Agilent 5977B MS. With a column flow of 0.9 mL/min, the GC oven was heated from 85°C to 150°C at a rate of 12°C/min and held for 3.5 min. MS settings and data analysis were the same as previously described. Standard curves on the TD-GC-MS were generated by sampling varying volumes (10 to 400 mL) of a 21 ppb CH_3_Br, 510 ppb CH_3_Cl (mol/mol) standard (10% analytical uncertainty, balance N_2_, Airgas). A conversion factor (peak area/volume sampled/CH_3_X concentration) was calculated from the slope of the standard curve (peak area/volume sampled) and used to calculate CH_3_X concentration in experimental samples using Equation 1:

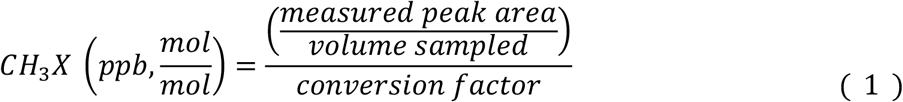

### Soil CH_3_X emissions

To assess headspace CH_3_X above soil samples lacking engineered microbes, twice-autoclaved or unsterilized B horizon soil (1 kg) was added to an incubation chamber and hydrated to 64% field capacity using MIDV1 medium (-N) with varying NaBr (0 to 100 mM). To establish the minimum number of engineered microbes required to yield an indicator gas signal, MG1655-*mht* cultures were grown to mid-log phase in M63 lacking halides, washed thrice, and resuspended in MIDV1 (-N) containing varying NaBr (0 to 100 mM). Unsterilized B horizon soil (1 kg) was then hydrated to 64% field capacity with this culture (200 mL) containing the specified cell population in a flask incubation chamber. For both experiments, flasks were sealed and incubated for 48 hours at 22°C prior to headspace TD-GC-MS analysis. Ambient samples were taken periodically inside our lab in Houston, TX between June 2021 and December 2021.

### IPTG sensing in soil

To see if soil affects IPTG bioavailability, MIDV1 (100 µL; -N, 20 mM NaBr) with varying concentrations of IPTG (0 to 1 mM) was added to empty GC vials and vials containing unsterilized B horizon soil (1 gram). Cultures of MG1655 transformed with pLC7 were grown to mid-log phase in M63 lacking halides and containing chloramphenicol (34 µg/mL). Cultures were washed thrice, resuspended in MIDV1 (-N; 20 mM NaBr) containing chloramphenicol (34 µg/mL), and added (100 µL) to the vials containing varying IPTG concentrations at a final cell population of 6×10^7^ CFU. Soil samples had a total water content of 64% field capacity. Samples were incubated at 22°C for 24 hours before headspace gas analysis. To benchmark indicator gas signals from the IPTG sensing strain against signals MG1655-*mht*, unsterilized B horizon soil (1 gram) was hydrated with MIDV1 (100 µL; -N, 100 mM NaBr) with or without IPTG (0 or 1 mM) in GC vials. Cultures of MG1655-*mht* and MG1655-IPTG were grown to mid-log phase in M63 lacking halides, with chloramphenicol (34 µg/mL) for the IPTG sensor only. Cells were washed thrice in MIDV1 (-N; 100 mM NaBr) without antibiotics, resuspended in the same medium, and added (100 µL) to the soil at a final cell density of 6×10^7^ CFU/gram; the total water content was 64% field capacity. Headspace gas was analyzed following a 24-hour incubation at 22°C. To investigate whether MG1655-IPTG presented a significant signal over background when added at a titer of 10^3^ CFU/gram soil, unsterilized B horizon soil (1 kg) was hydrated with MIDV1 (100 mL; -N, 100 mM NaBr) with or without IPTG (0 or 1 mM). Mid-log phase cultures of MG1655-IPTG in M63 lacking halides and containing chloramphenicol (34 µg/mL) were washed thrice and resuspended in MIDV1 (-N, 100 mM NaBr). Cultures (100 mL) were added to the pre-hydrated soils at a final density of 10^3^ CFU/gram soil and final water content of 64% field capacity. Sealed flasks were incubated at 22°C for 48 hours before gas analysis.

## SUPPORTING INFORMATION

**Supplementary Tables**: (S1) Soil particle size distributions, (S2) soil mineralogy, and (S3) plasmids used in this study.

**Supplementary Figures**: (S1) Soil XRD spectra, (S2) soil water retention curves, (S3) cell and halide titrations, (S4) *E. coli* survival in liquid medium, (S5) CH_3_Br standard persistence, (S6) *E. coli* survival in soil, (S7) TD-GC-MS methodology, and (S8) standard curves and ambient CH_3_X.

**Supplementary Methods**: (i) Materials and strains, (ii) soil particle size distribution, (iii) soil mineralogy, (iv) soil water retention curves, (v) indicator gas stability and cell viability, (vi) cell viability in untreated soil, and (vii) comparing GC-MS and TD-GC-MS measurements.

## ACKNOWLEDGEMENTS

We are grateful for financial support from the Defense Advanced Research Projects Agency HR0011-19-2-0019 (to C.A.M. and J.J.S.) and Gordon and Betty Moore Foundation (to C.A.M. and J.J.S.). Additionally, this research was supported by the US Department of Energy, Office of Science, through the Genomic Science Program, Office of Biological and Environmental Research, under FWP 78814 at PNNL. PNNL is a multi-program national laboratory operated by Battelle for the DOE under Contract DE-AC05-76RLO 1830. We would like to thank Imna Melendez, Jennifer Vu, Cassia Lewandowski, and Angela Sanchez for their technical support.

