## Supplemental Information for "Non-destructive chemical sensing within bulk soil using 1000 biosensors per gram of matrix"

### LIST OF SUPPLEMENTAL DATA


**Supplemental Tables**

**Table S1.** Soil particle size distribution.

**Table S2.** Soil Mineralogy.

**Table S3**. Plasmids used in this study.

**Supplemental Figures**

**Figure S1.** Soil XRD spectra.

**Figure S2.** Soil water retention curves.

**Figure S3.** Cell and halide titrations.

**Figure S4.** *E. coli* survival in liquid medium.

**Figure S5.** $CH_3Br$ standard persistence.

**Figure S6.** *E. coli* survival in soil

**Figure S7.** TD-GC-MS methodology.

**Figure S8.** Standard curves and ambient $CH_3X$ variability.

**Supplemental Methods**

**Materials and strains.**

**Soil particle distribution.**

**Soil mineralogy.**

**Water retention curves.**

**Indicator gas stability and cell viability.**

**Cell viability in untreated soil.**

**Comparing GC-MS and TD-GC-MS measurements.**

**Table S1.** Particle size distribution of the Texas forest Alfisol. Data represent the average of three technical replicates (±1 standard deviation).

| Horizon | Sand (%) | Silt (%) | Clay (%) |
|---|---|---|---|
| A (0-10 cm) | 77.7 (±1.0) | 8.7 (±0.1) | 13.6 (±0.5) |
| B (10-58 cm) | 80.2 (±0.6) | 11.8 (±0.5) | 8.1 (±0.8) |

**Table S2.** Soil mineralogy of the Texas forest Alfisol. Data represent the average of three technical replicates (±1 standard deviation).

| **Horizon (Depth)** | **Quartz (%)** | **Albite (Na feldspar) (%)** | **Kaolinite (%)** |
|---|---|---|---|
| A (0-10 cm) | 77.0 (±9.5) | 14.2 (±10.0) | 8.8 (±1.1) |
| B (10-58 cm) | 80.2 (±8.0) | 16.0 (±6.5) | 3.8 (±0.8) |

**Table S3.** Constructs used in this study.

| Plasmid | Description | Promoter (Regulation) | ORF | Addgene ID |
|---|---|---|---|---|
| pEMF051 | Constitutive sfGFP-MHT expression | P14-BCD22 (Constitutive)[1] | sfGFP-$(GGGGS)_2$-MHT | 182161 |
| pLC7 | IPTG-regulated MHT expression | $P_{LacO-1}$ (LacI)[2] | MHT | 182162 |
| pBAV1K-T5-gfp[3] | GFP expression with kanamycin resistance | T5 (LacI) | GFP | 26702 |

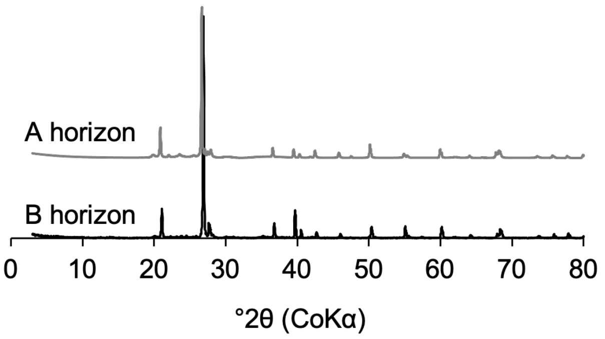


**Figure S1. XRD spectra of the Texas forest Alfisol.** XRD spectra for A (gray line) and B (black line) horizons.

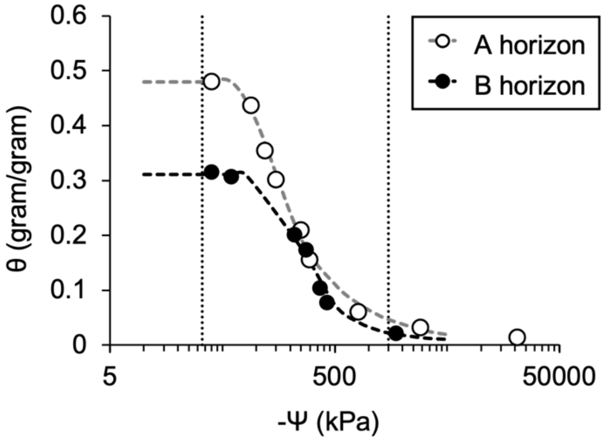


**Figure S2. Water retention curves of the Texas forest Alfisol.** Measured total soil potential (-Ψ) at varying water content (θ) (gram $H_2O$/gram dry soil). Data represent experimental measurements for the A (○) and B (●) horizons, while the dashed line represents empirical model predictions for the A (gray line) and B (black line) horizons using the van Genuchten model[4,5]. Field capacity (-33 kPa) and permanent wilting point (-1500 kPa) are represented as vertical lines.

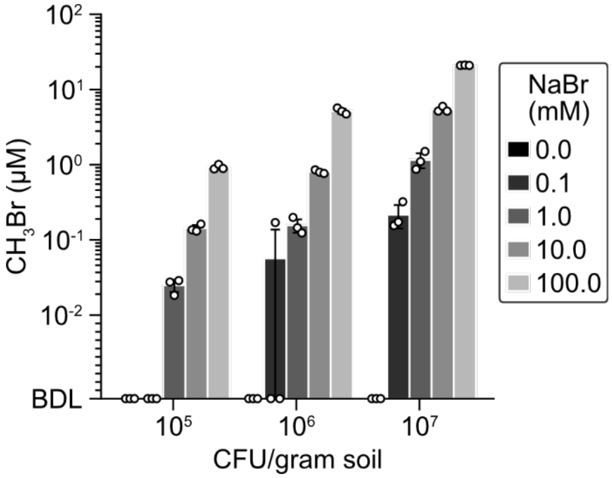


**Figure S3. Indicator gas measurements at varying cell titers and halides.** $CH_3Br$ signal presented by MG1655-*mht* ($10^5$ to $10^7$ CFU/gram soil) in twice-autoclaved B horizon soil (1 gram) hydrated to 64% field capacity with MIDV1 (-N; 0 to 100 mM NaBr). Samples were incubated for 24 hours at 22°C before headspace gas analysis. Bars represent the average of three biological replicates, while open circles represent individual data points. Error bars represent ±1 standard deviation. BDL, below detection limit.

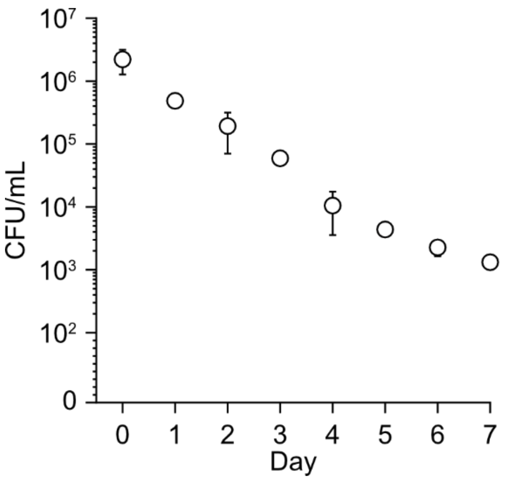


**Figure S4. Cell viability in liquid culture.** MG1655 cells expressing a GFP-MHT fusion protein were incubated in M63 medium (1 mL; -N, 100 mM NaBr) for seven days at 37°C with 250 rpm shaking. Matched samples were sacrificed each day, and serial dilutions of each culture were plated on LB-agar with chloramphenicol (34 µg/mL). The plates were incubated overnight 37°C and to enable CFU counting. Data represent the average of three biological replicates, while error bars represent ±1 standard deviation.

**a**

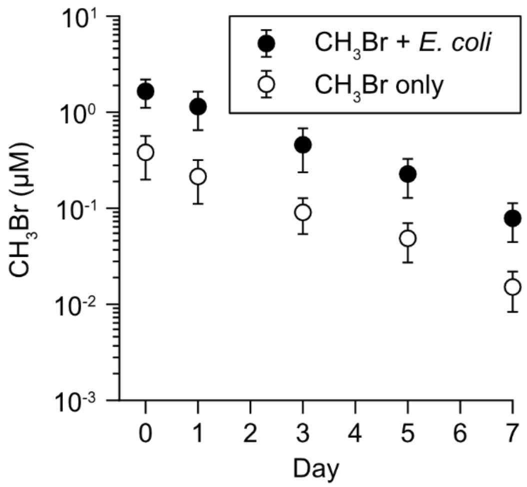


**b**

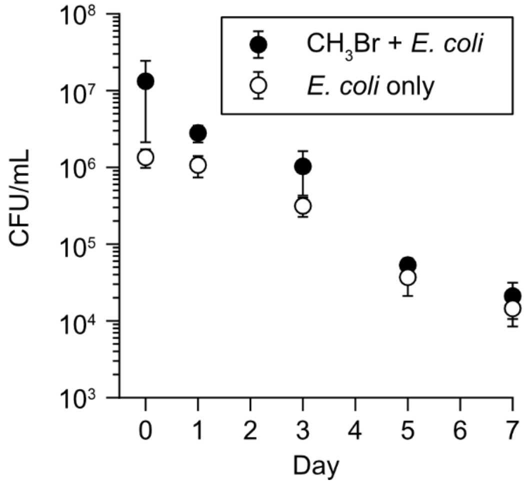


**Figure S5. $CH_3Br$ standards and cell viability over 7 days. (a)** Indicator gas measurements from a $CH_3Br$ standard incubated at 37°C, 250 rpm shaking for seven days. The standard was diluted in M63 (1 mL; -N, 100 mM NaBr) with (●) or without (○) MG1655 ($10^5$ CFU/mL). Samples were sacrificed every 24 hours for headspace gas analysis. **(b)** Viable MG1655 cells incubated in M63 (1 mL; -N, 100 mM NaBr) with (●) or without (○) $CH_3Br$ (0.02 µM) for seven days at 37°C, 250 rpm shaking. Samples were sacrificed every 24 hours, serial dilutions of each culture were plated on LB-agar, and plates incubated overnight at 37°C to enable CFU counting. Data represent the average of three biological replicates, and error bars represent ±1 standard deviation.

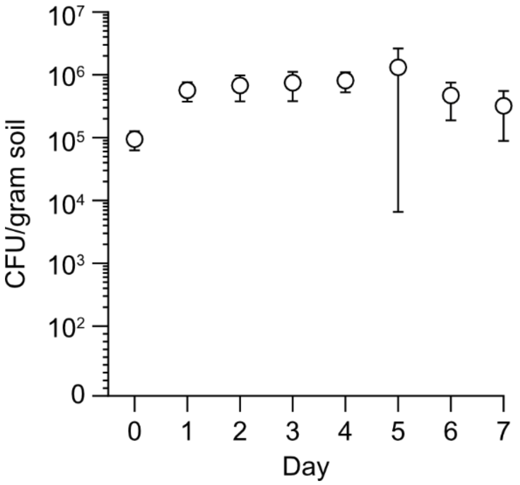


**Figure S6. *E. coli* viability in untreated soil.** MG1655-*mht* cells carrying a plasmid with a kanamycin resistance marker were suspended in MIDV1 (-N; 20 mM NaBr) with kanamycin (50 µg/mL). Untreated B horizon soil (1 gram) was hydrated to 64% field capacity with culture ($6x10^7$ CFU/gram soil) and incubated at 22°C in sealed vials. At each timepoint, samples were sacrificed, and half of the hydrated soil (0.6 gram) was vortexed for 10 minutes in 1 mL sterile PBS buffer. The soil particles were allowed to settle, and serial dilutions of the supernatant were plated on LB-agar plates with kanamycin (50 µg/mL) and incubated overnight at 37°C to enable CFU counting. Data represent the average of three biological replicates, while error bars represent ±1 standard deviation.

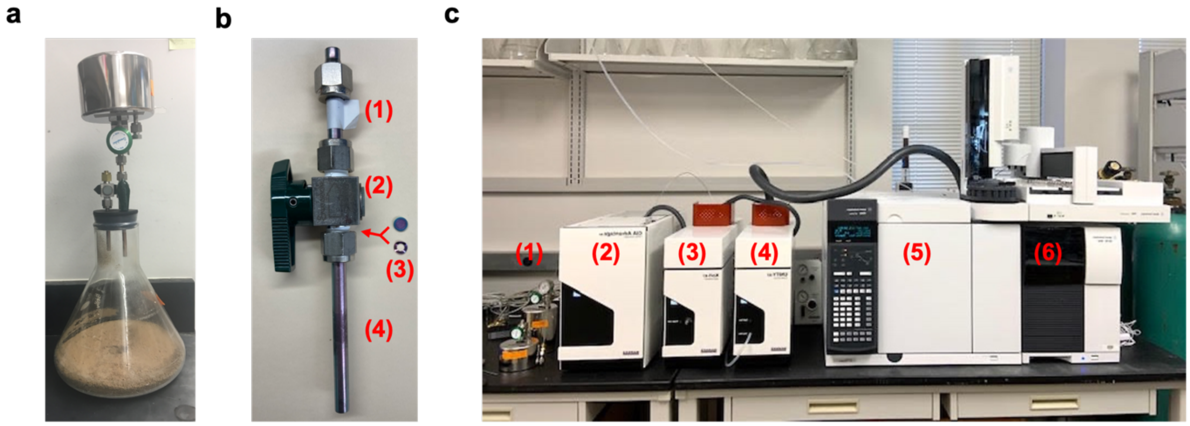


**Figure S7. TD-GC-MS sampling equipment. (a)** Flask chamber for incubation experiments. A 2.8 L Fernbach flask is closed with a two-holed rubber stopper fit with two sets of valved sampling tubes. An evacuated 0.5 L canister is connected to enable headspace sampling. **(b)** A sampling tube assembly, consisting of (1) ¼" tubing for attaching the canister, (2) a plug valve, (3) a 0.2 μM frit filter and washer (inside the tube fitting), and (4) ¼" tubing to insert into the stopper. **(c)** TD-GC-MS equipment, consisting of: (1) sample canisters, (2) a canister autosampler, (3) a water removal unit, (4) a thermal desorption unit, (5) a gas chromatograph, and (6) a mass spectrometer.

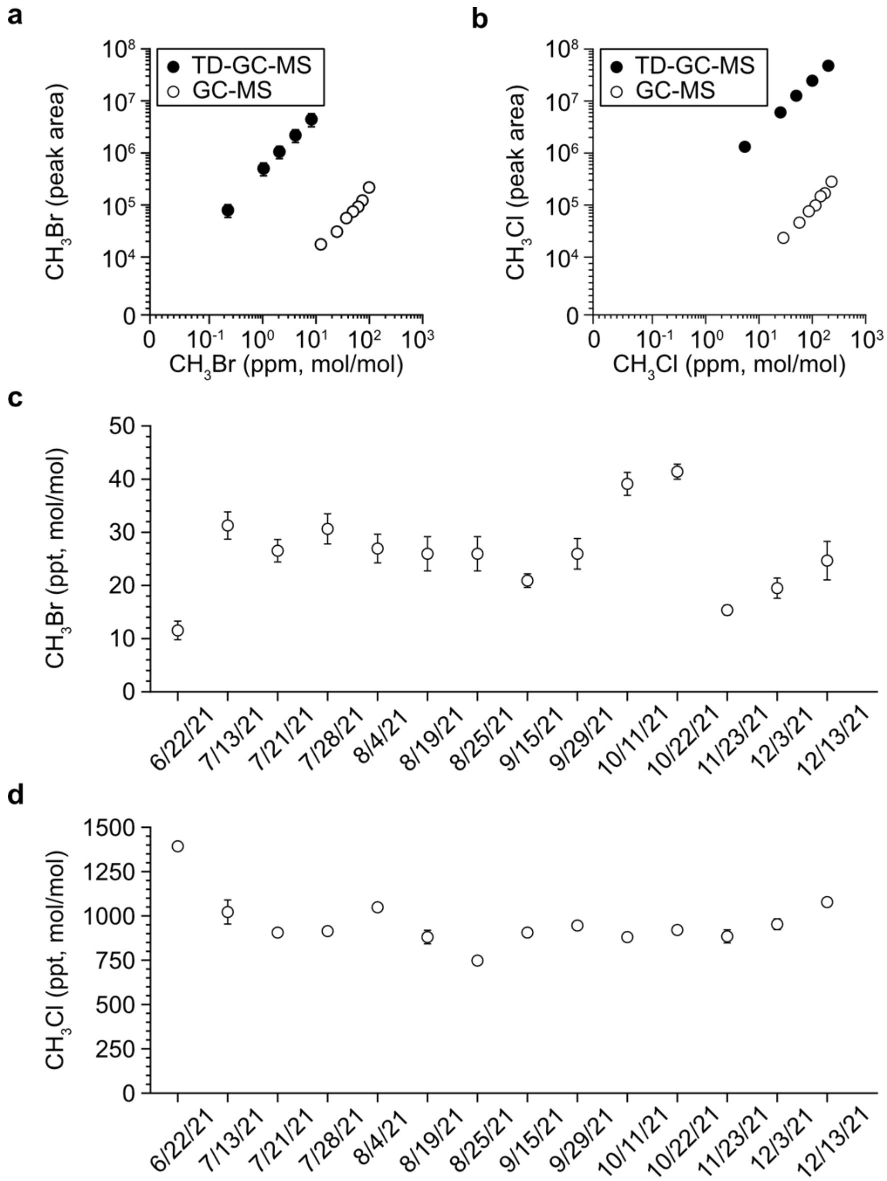


**Figure S8. Standard curves and ambient $CH_3X$ variability. (a)** $CH_3Br$ and **(b)** $CH_3Cl$ standard curves measured by TD-GC-MS (●) or GC-MS (○). For GC-MS measurements, liquid-phase standard curves were converted to mixing ratio by estimating gas partitioning using Henry's Law. For TD-GC-MS measurements, varying volumes of a single gas-phase standard were sampled, the total amount of $CH_3X$ in each sample was calculated, and the total $CH_3X$ for each sample was normalized to a standard volume (100 mL) to calculate the equivalent atmospheric mixing ratio of samples with varying $CH_3Br$ concentrations. The comparative peak area from TD-GC-MS to GC-MS samples represents the change in sensitivity to $CH_3X$. **(c)** Ambient $CH_3Br$ and **(d)** ambient $CH_3Cl$ measured indoors at Rice University in Houston, TX. Data represent the average of three independent replicates, while error bars represent ±1 standard deviation.

### SUPPLEMENTARY METHODS

**Materials and strains.** Kanamycin was from Research Products International. To enable selection of MHT-expressing cells from soil samples, a plasmid carrying a kanamycin resistance marker (pBAV1K-T5-gfp)[3] was transformed into MG1655-*mht*. pBAV1K-T5-gfp was a gift from Ichiro Matsumura (Addgene plasmid #26702; http://n2t.net/addgene:26702; RRID: Addgene_26702). All other materials are as described in the manuscript.

**Soil particle distribution.** Soil particle size distribution was measured using chemical dispersion followed by gravity sedimentation[6]. Soils (30 g) were suspended in 3% hexametaphosphate (HMP) solution in a 250 mL high density polyethylene (HDPE) bottle with 3:1 HMP (90 mL) to soil (30 g) ratio. The suspension was mixed on an orbital shaker (OS-500, VWR) for 2 hours. The sand fraction was then collected using a USA No. 270 standard test sieve (53 µm). The remaining silt and clay suspension fraction was stirred thoroughly and allowed to settle undisturbed in a 200 ml volumetric cylinder for a sedimentation period of 90 min. After the sedimentation, the suspended clay fraction was decanted. The sand fraction collected from sieving and the settled silt fraction was then dried in an Al weighing dish (20 mL; HS14521A, Heathrow Scientific) at 60 °C to constant weight. The clay fraction was calculated by subtracting the weight of sand and silt from the original sample mass.

**Soil mineralogy.** Bulk powder diffraction patterns were obtained using a X-ray diffractometer (D/MAX 2100, Rigaku) with Cu K-alpha radiation and a graphite monochromator operated at 40 KeV and 40 mA. Aliquots consisting of ~0.5 g soil were finely ground and homogenized in an agate mortar and pressed powder mounts were prepared in 15 × 20 mm Al sample holders. The samples were scanned from 3 to 80° 2θ at 0.02° steps with 60 second measurement time per step. Data analysis was performed using Rigaku PDXL2 software.

**Water retention curves.** Soil samples were rewetted with deionized water to take the water content from 0 gram $H_2O$/gram dry soil to a value near the field capacity. Both the dry and wet mass of the soil sample were recorded with an analytical balance and used to calculate the water content (θ). About 7.5 ml of each sample was then placed into a 15 ml stainless steel sample cup (Decagon Devices Inc., Pullman, WA). The chamber was covered with a plastic cap for 2-3 hours to allow moisture equilibration across the whole chamber. The plastic cap was then removed, the cup was inserted into the sample chamber, and the soil potential (ψ) by a dewpoint potentiometer (WPC4, Decagon Devices) at 23°C. The measured data was then fitted using SWRC Fit[5] with the van Genuchten model[4].

**Indicator gas stability and cell viability.** To assess $CH_3Br$ signal persistence over one week with or without *E. coli*, $CH_3Br$ (0.02 µM) was added to M63 (1 mL; -N, 100 mM NaBr), with or without MG1655 cells (6x$10^4$ CFU/mL) in GC vials. Samples were sealed, incubated at 37°C with 250 rpm shaking. After 4 hours (day 0) and every 24 hours thereafter, matched samples were sacrificed for headspace gas analysis. To assess cell viability, samples of MG1655 (6x$10^4$ CFU/mL) were prepared and incubated as above with or without $CH_3Br$ (0.02 µM). At each timepoint, aliquots of a subset of samples were plated on LB-agar, and plates were incubated overnight at 37°C to enable CFU counting.

**Cell viability in untreated soil.** MG1655-*mht* carrying a plasmid with a kanamycin resistant marker (pBAV1K-T5-gfp) was grown to mid-log phase in M63 lacking halides and containing kanamycin (50 µg/mL). Cultures were washed thrice and resuspended in MIDV1 (-N; 20 mM NaBr) with kanamycin (50 µg/mL). Untreated B horizon soil (1 gram) was hydrated to 64% field capacity with culture (200 µL) to a final cell density of $6x10^7$ CFU/gram soil. Vials were capped and incubated at 22°C. Samples were sacrificed every 24 hours, and half of the hydrated soil (0.6 gram) was vortexed in PBS buffer (1 mL) for 10 minutes. Serial dilutions of the supernatant were plated on LB-agar plates with kanamycin (50 µg/mL) and incubated overnight at 37°C to enable CFU counting.

**Comparing GC-MS and TD-GC-MS measurements.** To compare the sensitivity of GC-MS to the sensitivity of TD-GC-MS, $CH_3X$ signals from liquid-phase standard curves were measured at 22°C. The concentration of $CH_3X$ in the headspace was estimated using Henry's Law (Equation S1):

$$C_{gas} = \frac{C_{liq}}{H^{cc} + \beta} \tag{S1}$$

where $C_{gas}$ is the concentration of $CH_3X$ in the gas phase (µM), $C_{liq}$ is the concentration of $CH_3X$ in the liquid phase (µM), $H^{cc}$ is the dimensionless solubility constant at 20°C ($CH_3Br$ = 4.214; $CH_3Cl$ = 3.223), and β is the ratio of headspace volume to liquid volume[7,8]. The gas-phase concentration was then converted to mixing ratio by assuming the ideal gas law for 1 mL of air in the vial headspace at 20°C and calculating the ratio of moles $CH_3X$/moles air.

For TD-GC-MS measurements, varying volumes (10 mL to 400 mL) of a 21 ppb $CH_3Br$, 510 ppb $CH_3Cl$ (mol/mol) gas-phase standard (10% analytical uncertainty, balance $N_2$, Airgas) were sampled from an 0.5 L canister. The total moles of $CH_3X$ in each sample was estimated by assuming ideal gas behavior at 20°C. The total moles of $CH_3X$ in each sample was normalized to a standard sample representative of TD measurements

(100 mL), and the mixing ratio of $CH_3X$ was calculated by assuming the ideal gas law for air. These approximations enabled the GC-MS and TD-GC-MS measurements to be directly compared under conditions representative of each sampling method.
